# Single cell RNA-sequencing reveals cellular heterogeneity and trajectories of lineage specification during murine embryonic limb development

**DOI:** 10.1101/659656

**Authors:** Natalie H. Kelly, Nguyen P.T. Huynh, Farshid Guilak

## Abstract

The coordinated spatial and temporal regulation of gene expression in the murine hindlimb determines the identity of mesenchymal progenitors and the development of diversity of musculoskeletal tissues they form. Hindlimb development has historically been studied with lineage tracing of individual genes selected *a priori*, or at the bulk tissue level, which does not allow for the determination of single cell transcriptional programs yielding mature cell types and tissues. To identify the cellular trajectories of lineage specification during limb bud development, we used single cell mRNA sequencing (scRNA-seq) to profile the developing murine hindlimb between embryonic days (E)11.5-E18.5. We found cell type heterogeneity at all time points, and the expected cell types that form the mouse hindlimb. In addition, we used RNA fluorescence in situ hybridization (FISH) to examine the spatial locations of cell types and cell trajectories to understand the ancestral continuum of cell maturation. This data provides a resource for the transcriptional program of hindlimb development that will support future studies of musculoskeletal development and generate hypotheses for tissue regeneration.

## Introduction

The development of the mammalian limb, and subsequently the synovial joints, involves a complex and coordinated sequence of events whereby a relatively homogeneous limb bud differentiates into multiple cell types forming the tissues in the limb. While certain aspects of limb development have been deciphered, a more thorough understanding of joint development may provide important insights into the development of methods to enhance regeneration of musculoskeletal tissues.

Mapping of the precise sequence of events involved in limb development may provide new targets to stimulate regeneration of musculoskeletal tissues. Although limb development has been studied via the expression of pre-determined markers and fate-mapping, the trajectories of gene expression that drive individual cell differentiation toward distinct lineages are not fully understood due to limitations in *a priori* selection of limited lineage markers and the effects of combining all cells for conventional “bulk” transcriptomic analyses.

In this regard, advances in single cell gene expression analysis have made it possible to decipher cell transcription at increasingly high resolution. Massively parallel droplet based assays can determine the transcriptional profiles of thousands of single cells simultaneously [1-3]. The availability of these new tools has led to an increasing number of initiatives to create comprehensive cell atlases in the human [4, 5], and mouse [6, 7]. These cell atlases focus on adult tissues, while recent work in the mouse has examined gastrulation and organogenesis of whole organisms at single-cell resolution [8, 9]. Other studies have focused on particular organ or cell-type development, revealing previously unappreciated heterogeneity in neuronal, myocardial, and pulmonary development [10-14]. Recently, a single-cell transcriptomic atlas of limb development in the chick highlighted the transcriptional complexity of the 23 distinct cell populations in the autopod [15].

These studies have enhanced the understanding of development of these tissues and provided resources and data to advance regenerative medicine. However, such single-cell data has not been reported for the development of the mammalian limb. The goal of this study was to quantify the transcriptional landscape during mouse limb development via single cell RNA-sequencing and to determine the trajectories of cellular lineage specification into joint tissues during this process.

## Results

To examine the transcriptional landscape of murine hindlimb development, the limb bud was isolated at four developmental time points: embryonic day 11.5 (E11.5) to examine early limb bud formation, E13.5 when cartilaginous condensations form, E15.5 for joint cavitation, and E18.5 when limb morphogenesis has progressed. Single cells were dissociated from hindlimb tissues and droplet-based high-throughput single-cell RNA-sequencing was performed (10x Genomics) (Fig 1a) [3]. In addition, a species mixing experiment of mouse E18.5 hindlimb cells combined 1:1 with human 293T cells from culture was analyzed to determined single cell capture efficiency. Single cell capture was highly reliable as demonstrated by a low (1.4%) multiplet rate (Supplemental Figure 1). mRNA from an average of 2,500 cells per experimental time point was captured and sequenced with an average of 200K sequencing reads per cell. We detected the expression of an average of 2,700 genes per cell in each sample.

**Figure 1:**
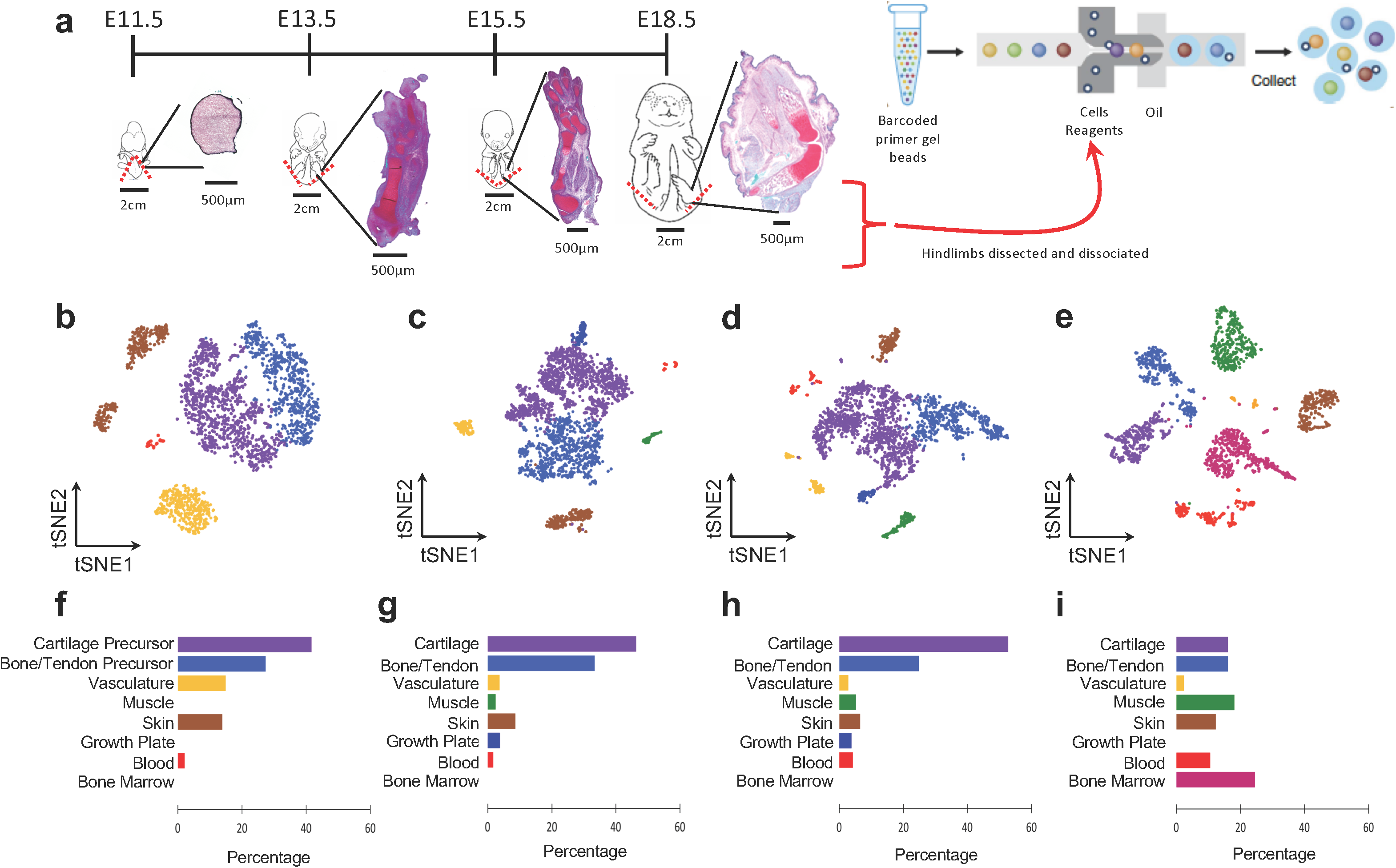
Distinct populations can be determined in mouse embryonic hindlimb cells. **(a)** Schematic of sampling area at the four stages of hindlimb development (whole embryo schematics adapted from EMAP eMouse Atlas Project http://www.emouseatlas.org [30]). Both hindlimbs were dissected (indicated by red dashed lines) from n=6-7 pups per litter. Insets demonstrate cartilaginous morphology at each time point via Safranin-O/Fast Green/Hematoxylin staining. Hindlimbs were dissociated to single cells and captured using 10x single cell RNA-seq technology. **(b-e)** tSNE projection of the four developmental time points, containing 2,627 (E11.5), 2,815 (E13.5), 2,596 (E15.5) and 1,901 (E18.5) cells, where each cell is grouped into clusters (distinguished by their colors). Comparable cell populations identified in multiple samples are visualized using the same color. **(f-i)** Percentage of cells represented by each cluster. Colors correspond to the clusters identified in **(b-e)**.

Cluster analysis within each time point yielded between 5-7 distinct clusters which separated across time (Fig 1b-e) [16]. At E11.5, we identified 5 cell types representing cartilage precursors (*Hoxd13* expression), bone/tendon precursors (*Tbx13*), skin (*Krt14*), vasculature (*Cdh5*), and blood (*Lyz2*) (Fig 1b,f and Supplemental Fig 2a). At E13.5, cartilage and bone/tendon clusters expressed more mature marker genes (*Col2a1* and *Col1a1*, respectively). In addition to skin, vasculature, and blood clusters there were additionally clusters representing the growth plate (*Col10a1*) and muscle (*Myog*) at E13.5 (Fig 1c, g and Supplemental Fig 2b). At E15.5, the number of clusters and cell type percentages did not differ from E13.5 (Fig 1d,h and Supplemental Fig 2c). At both E13.5 and E15.5 the majority of cells were cartilaginous. Finally, at E18.5 there was no longer a cluster representing the growth plate, and cell type percentages were more evenly distributed (Fig 1e,i and Supplemental Fig 1d).

**Figure 2:**
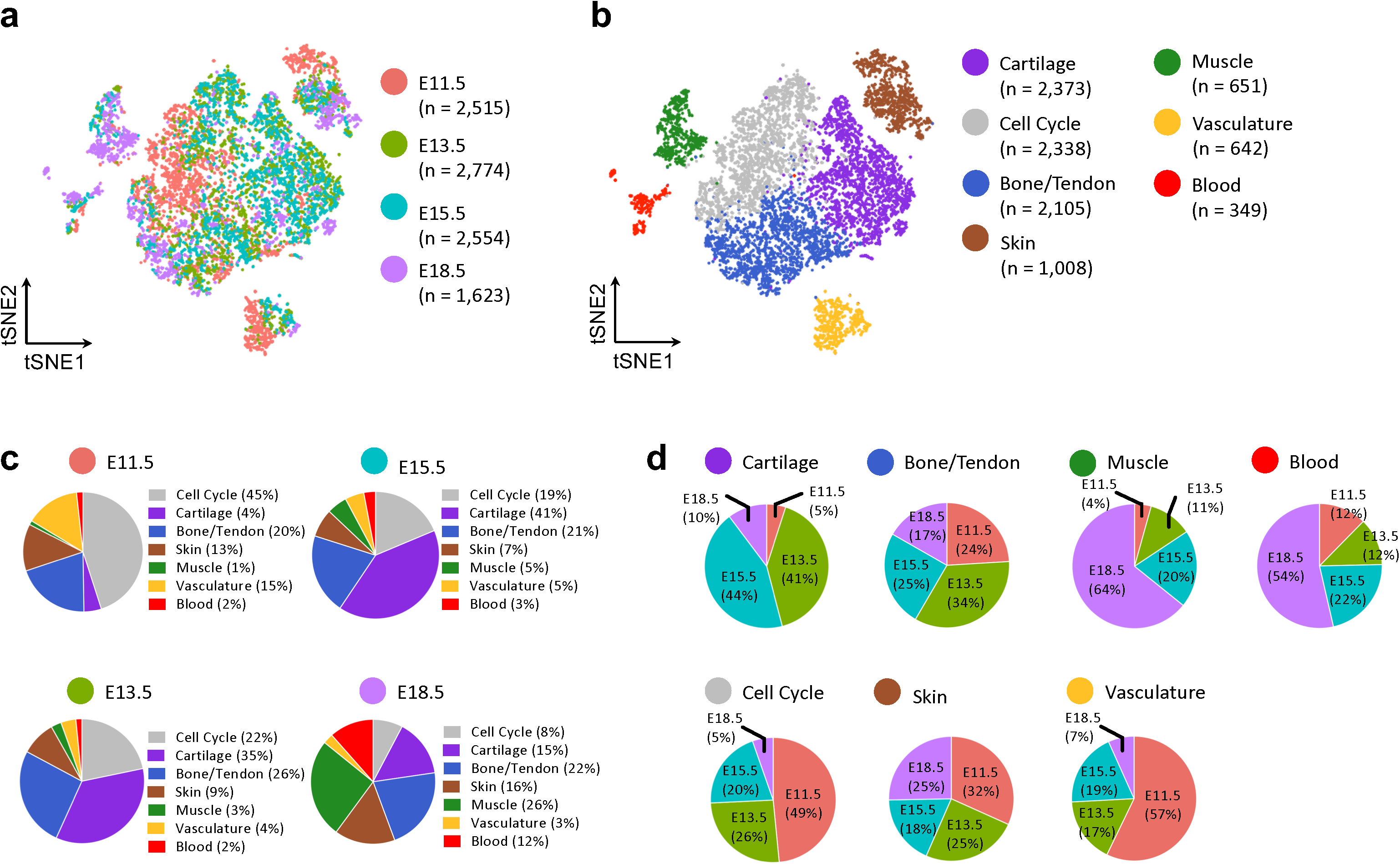
tSNE projection of the merged dataset with cells colored by **(a)** time point and **(b)** cell type determined by unsupervised clustering. **(c)** Percentage of cell type at each time point and **(d)** percentage of cells in each type by time point.

To determine the similarity of cell types across time points, canonical correlation analysis was performed [17]. Across all time points there were 9,466 cells in 7 clusters, with each cluster containing cells from all time points (Fig 2). Even following cell cycle regression, a cluster emerged with gene expression enriched for mitotic nuclear division (GO:0140014). Cells in this cluster were primarily from the earliest time point (E11.5, 49%), while at E18.5 only 8% of cells were assigned to the cell cycle cluster. The six other cell type clusters recapitulated the cell types found in the individual time points: cartilage, bone/tendon, skin, muscle, blood, and vasculature. The cartilage cluster was mostly comprised of cells from the middle time points (E13.5, 41% and E15.5, 44%), reflecting the prevalence of cartilaginous skeletal elements. As seen in the individual time point clustering, there were very few cells at E11.5 (4%) that were clustered as muscle. In fact, the muscle cluster was comprised mostly of cells from E18.5 (64%). Likewise, the blood cluster was mostly E18.5 (54%).

To examine developmental trajectories we used an unsupervised algorithm, p-Creode, that produces multi-branching graphs from single-cell data [18]. E11.5 and E18.5 cells were found at the ends of the cell trajectory, with E13.5/E15.5 cells along the middle of the trajectory branches (Fig 3a). A large branch consisting of E11.5 cells was comprised of musculoskeletal precursors which gave rise to cartilage (Fig 3a, purple arrow; Fig 3b *Col2a1* expression overlay), and bone/tendon (Fig 3a, blue arrow; Fig 3b *Col1a1* expression overlay). While muscle cells arose from later cells (E13.5) along this same branch (Fig 3a, green arrow; Fig 3b *Myog* expression overlay). A separate branch defined the developmental trajectory of skin (Fig3a, brown arrow; Fig 3b *Krt14* expression overlay). Lastly, another branch expressed both vasculature (Fig 3b *Cdh5* expression overlay) and blood (Fig 3a, red arrow; Fig 3b *Lyz2* expression overlay) markers.

**Figure 3:**
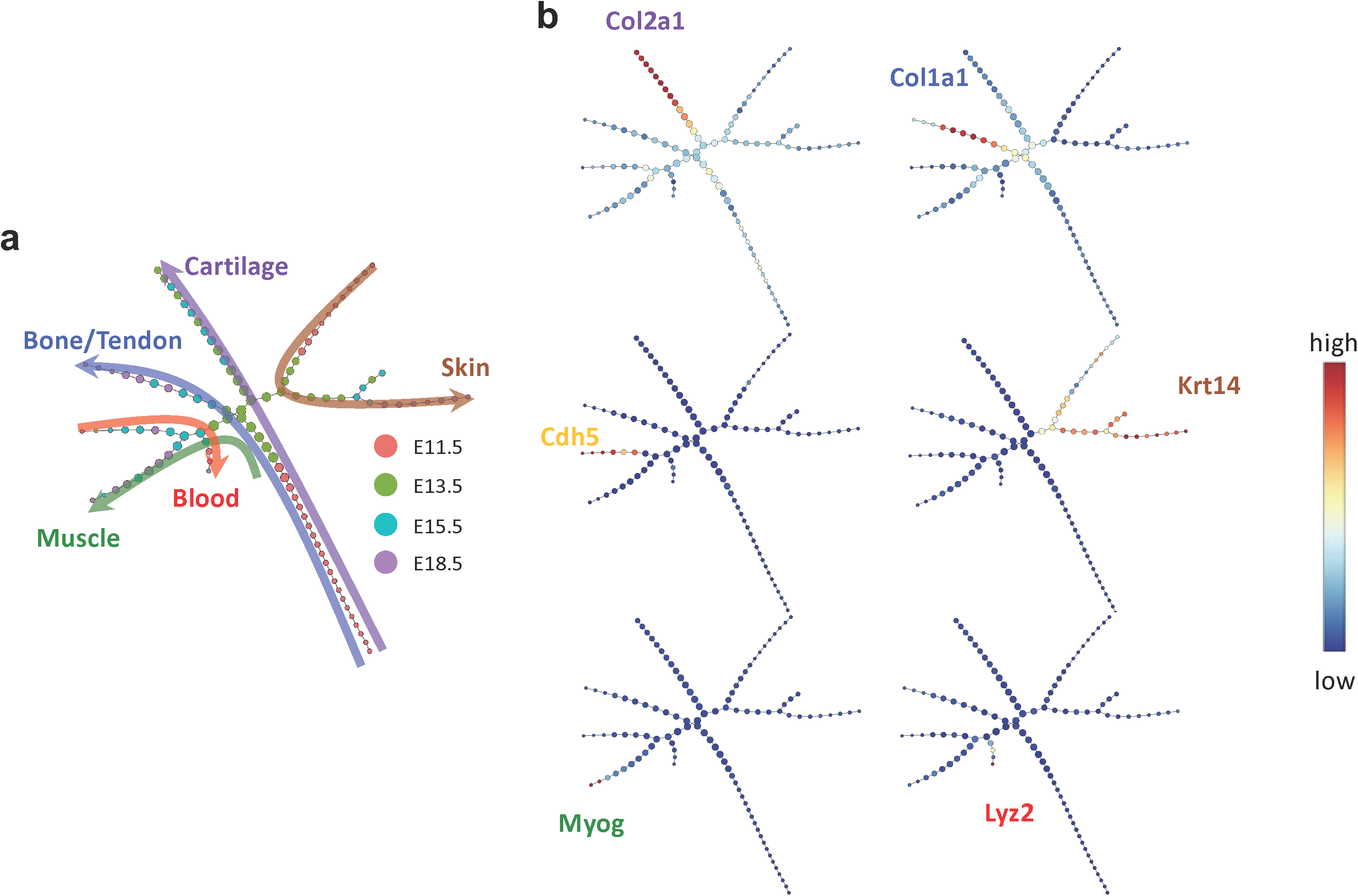
(a) p-Creode trajectory analysis with node colors indicating time point and colored arrows indicating tissue type across developmental trajectory. **(b)** Overlay of selected transcripts depicting cartilage (Col2a1), bone/tendon (Col1a1), vasculature (Cdh5), skin (Krt14), muscle (Myog), and blood (Lyz2) development in the hindlimb on the pCreode topology generated in **(a)**.

To verify the spatial distribution of specific marker genes and time point expression profiles, we performed RNA FISH for two sets of probes. One set examined the expression of a cartilage marker gene (*Col2a1*), a skin marker gene (*Krt14*), and a muscle marker gene (*Myog*). We found that cartilage and skin were evident at all time points, while muscle was not visible until the latest time point (E18.5). *Col2a1* was expressed across a diffuse area of the hindlimb bud at E11.5, which coalesced into cartilaginous rays indicating the future sites of bones in the hind paw at E13.5, while at E15.5 and E18.5 the cartilaginous rays were now interrupted, indicating joint cavitation had occurred. Furthermore, at E18.5, *Col2a1* expression was further restricted to the joint surfaces (Fig 4).

**Figure 4.**
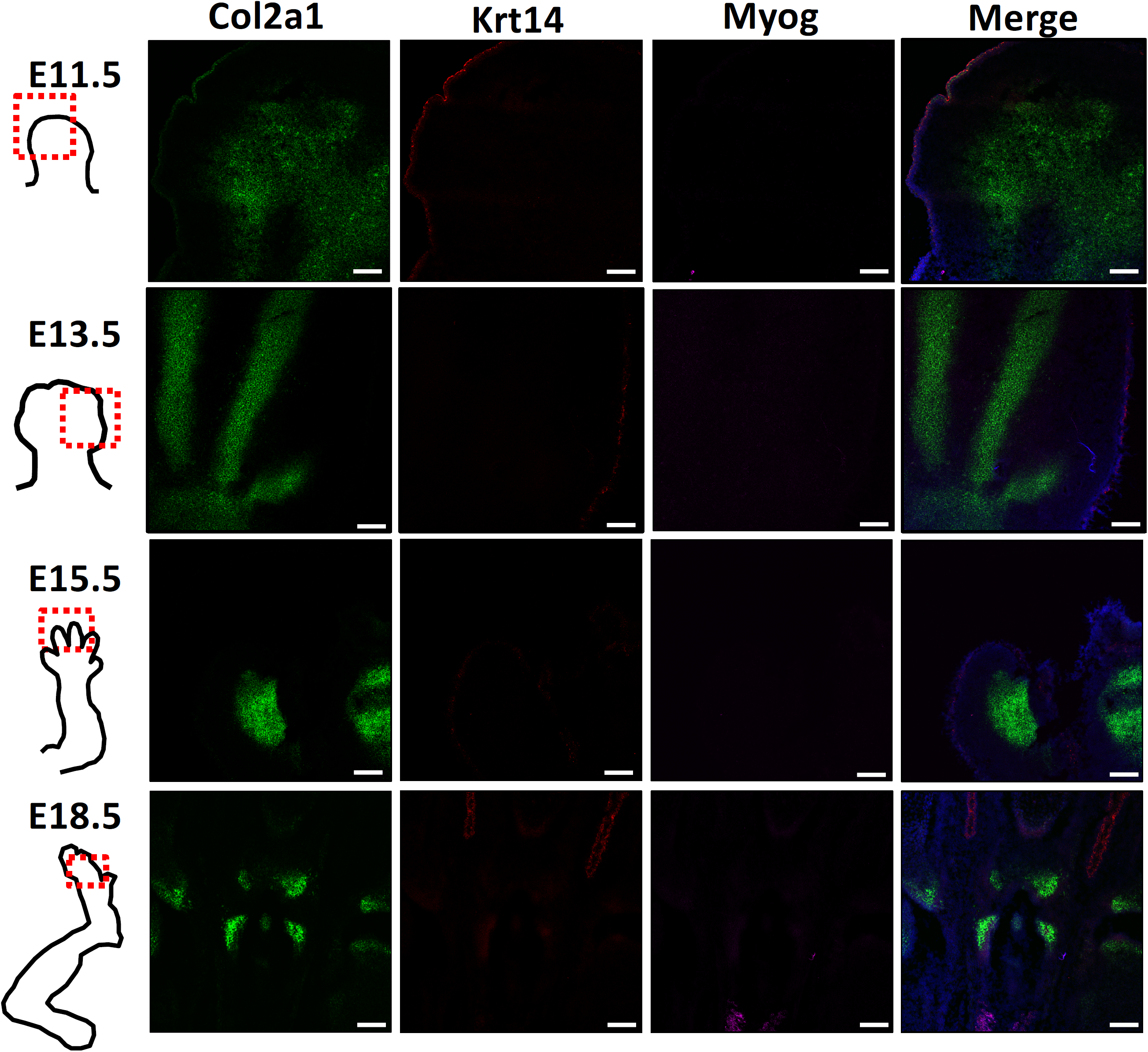
Representative RNA FISH images of hindlimbs at four developmental time points to verify spatial locations of cartilage (Col2a1), skin (Krt14), and muscle (Myog). Scale bar = 100 μm

The second set of marker probes were selected to better understand the cell clusters determined via tSNE that were marked by *Col1a1* expression. Because we did not identify a specific cell cluster at any time point marked by tendon gene expression (e.g. Scleraxis, *Scx*), we wanted to spatially verify the expression of *Col1a1, Scx* (to identify tendon), and *Runx2* (to identify bone). At E11.5 there was not clear expression of the three markers. At E13.5 there was faint staining of *Col1a1*, along the rays of the cartilaginous condensations. By E15.5 *Runx2* was found along the nascent bone, both overlapping and, separate from, *Col1a1*. Minimal *Scx* was seen at E15.5, which did not overlap with *Col1a1* or *Runx2*. Finally, at E18.5 bone and skin were marked by *Runx2* expression, while *Col1a1* was in locations on bone that overlapped with *Runx2*, and along the sides of the bone, possibly marking tendon. However, *Scx* expression was not present (Fig 5).

**Figure 5.**
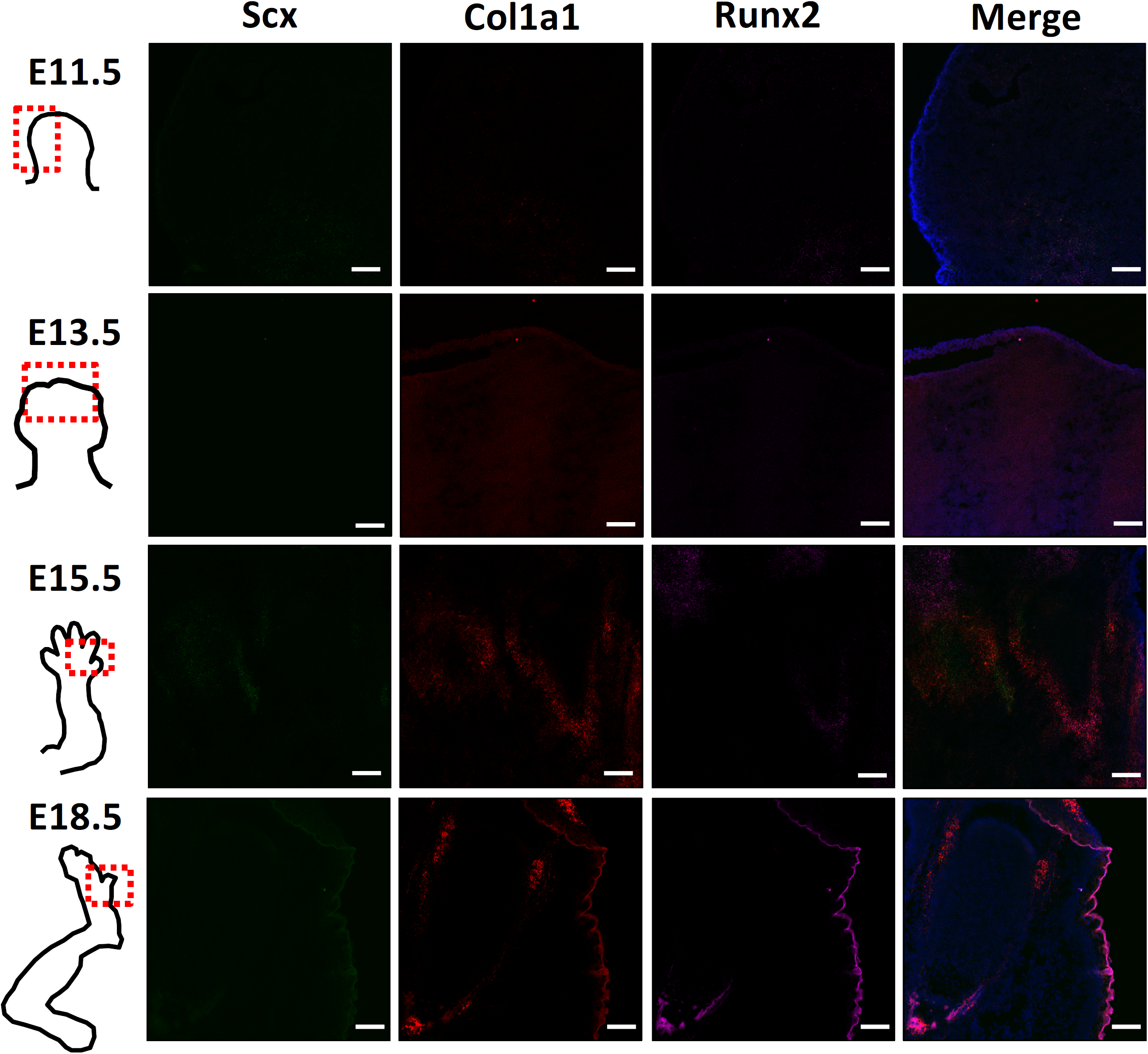
Representative RNA FISH images of hindlimbs at four developmental time points of tendon (Scx) and bone (Runx2) markers, as well as Col1a1, to verify what cell types are defined by clusters high in Col1a1. Scale bar = 100 μm

## Discussion

We successfully generated single cell RNA-sequencing data profiling the transcriptional landscape of murine limb development. We observed cell type heterogeneity even at the earliest developmental time point that a limb bud could be manually isolated (E11.5). The identified cell clusters recapitulated the known cell types present in hindlimb development, i.e., cartilage, bone, muscle, skin, vasculature and blood. Trajectories of musculoskeletal cells identified five main developmental branches that help to explain the ancestral continuum of cell maturation. These findings provide the basis for further understanding of the process of early limb and joint development, which will hopefully serve to inform new approaches for deconstructing and eventually recapitulating the regenerative process.

Cell type heterogeneity at the early limb bud stage (E11.5) was observed, with musculoskeletal precursors clustering into two populations which seem primed even at this early stage to develop into cartilage or bone/tendon. Similarly, five cell type clusters where found at a roughly equivalent developmental time point in the chick autopod (HH25, approximately mouse E12 [19]), although muscle cells were found in chick, but not mouse [15]. Genes indicating the presence of Muscle muscle cells were apparent at E13.5 by scRNA-seq, but were not observed with RNA FISH until E18.5. The lack of signal in RNA FISH until the latest time point could be explained by the single, 2D section. Other cell type clusters included blood, vasculature, bone, cartilage, and skin.

Hypertrophic chondrocytes are typically marked by *Col10a1* expression and are prevalent in the growth plate. We found clusters defined by *Col10a1* expression at the middle developmental time points, E13.5 and E15.5, but not at E18.5. This, again, may be due to cell type drop-out due to limitations in cell capture or it may be that hypertrophic chondrocytes are more similar to bone or cartilage cells at the most mature time point.

Spatial expression information is lost with scRNA-seq, as cells must be dissociated from the tissues. We used RNA FISH to examine the spatial expression of markers for cartilage, skin, and muscle. In addition, we used a second set of markers to determine the spatial location of cells with high expression of *Col1a1*, and to see if these cells were at locations of bone, tendon/ligament, or both. It appears that *Col1a1* may be expressed both at sites of bone (overlap with bone marker, *Runx2*), and possibly at sites of tendon/ligament as well (due to spatial location at edges of cartilage/bone surface).

Cells were aligned along developmental trajectories that defined cartilage, bone, skin, muscle, and blood. Future work will further explore the transitional gene expression that determines whether an early mesenchymal progenitor becomes cartilage or bone, for example. This information may be used in in vitro systems to more robustly convert embryonic, mesenchymal, or induced pluripotent cells to musculoskeletal tissues for tissue engineering applications. A well characterized marker of joint formation, *Gdf5*, was not highly expressed in the cells in our study. Gdf5 is a transient marker of the interzone, and is present in a restricted region that is only a few cell layers thick and is thus only expressed in a small number of cells [20, 21]. This could be due to limitations in cell capture or the need for larger cell numbers to robustly quantify rare cell populations. Furthermore, it was difficult to classify the clusters high in collagens type I and III as bone, tendon or ligament because there was not clear expression of tendon markers scleraxis (*Scx*) or tenomodulin (*Tnmd*) or bone markers *Runx2* or osteocalcin (*Bglap*). Deeper sequencing may be necessary to more accurately define this cell population and gain insight into transcripts that may have relatively low expression levels.

In conclusion, this work provides an in-depth resource describing the development of the mouse hindlimb. In combination with other genetic methods such as lineage tracing, this approach could provide new insights into the interaction among different cell types. A thorough understanding of the limb development process will hopefully allow recapitulation of these events to enhance joint repair or, eventually, joint or limb regeneration.

## Materials and methods

### Animals

Murine hindlimbs were manually dissected at 4 time points during embryonic joint development, including points marking formation of the limb bud (E11.5), formation of cartilaginous condensations (E13.5), joint cavitation (E15.5), and joint morphogenesis (E18.5). Embryonic pups (7-9/litter) were removed by caesarean section from a timed-pregnant female C57BL/6 mouse per time point (The Jackson Laboratory, Bar Harbor, ME) and euthanized. Hindlimbs from 6-7 pups/litter were digested with collagenase (Type II, Worthington-biochem, Lakewood, NJ) and pronase (EMD Millipore, Billerica, MA) in 15-minute increments with agitation at 37°C for up to one hour. Every 15 minutes mechanical dissociation was applied by pipetting up and down. Following digestion, cells from each litter (time point) were pooled, strained through a 100μm cell strainer, pelleted, resuspended at 1 × 10^6^ cells per 500μl in freeze media (80% FBS, 10% DMEM, 10% DMSO), and stored in liquid nitrogen until single cell capture. All experiments were approved by the Washington University IACUC.

### Single Cell Capture, and RNA-sequencing

Cells in freeze media were rapidly thawed in a 37°C water bath, and resuspended in 1X PBS and 0.04% bovine serum albumin. Dead cells were removed using magnetic bead-bound antibodies against apoptotic and necrotic cells (Dead cell removal kit, Miltenyi Biotec) and cells were washed twice. Single cell capture was performed with the 10x Chromium Controller (10x Genomics, Pleasanton, CA) [3]. Briefly, single cell suspensions were brought to a concentration of 1,000 cells/μl and approximately 6,000 cells per time point (4 samples total) were loaded into the 8-channel microfluidic chip that encapsulates thousands of cells in gel beads in emulsion (GEMs). Cell capture, cDNA generation, and library preparation were performed using Chromium Single Cell 3’ v2 Reagent kit, following the manufacturer’s instructions. To test the reliability of single-cell capture, a 1:1 mixture of E18.5 mouse hindlimb cells and human 293T cells was sequenced and the multiplet rate was determined by the fraction of mouse reads in human barcodes and vice versa. Single cell RNA-seq libraries were pooled and run on the Illumina NovaSeq S1 flow cell with 26x98bp reads to generate ∼500 million reads per library (time point).

### Data Analysis

Cell Ranger software (v2, 10x Genomics) was used to demultiplex samples, process barcodes, align to the mouse (GRCm38/mm10) or human (hg19; for mixed species) genome assembly, and count single cell genes [3]. The gene/barcode matrix was imported to Seurat (https://github.com/satijalab/seurat, v2.3.4) and filtering was performed to exclude cells with less than 200 or more than 6,000 genes. Cells with more than 10% unique molecular identifier (UMI) counts associated with mitochondrial genes were also excluded. The data was log-normalized and library size and mitochondrial UMI counts were regressed.

The cell cycle stage of each cell was assigned as S, G1, or G2/M based on a pre-determined list of cell cycle markers [22] and the difference between the G2/M and S scores was calculated and variance-corrected.

Cluster analysis was performed via t-distributed stochastic neighbor embedding (tSNE) [23] at each time point to examine cellular heterogeneity and cell types present in hindlimb tissues [17, 24]. In addition, the four time points were combined using Seurat’s Multi-canonical correlation analysis (CCA) command to identify projection vectors that maximize the overall correlation across all data sets [17, 24]. Cluster specific genes were determined by calculating the expression difference of each gene between that cluster and the average of the rest of the clusters. The top 100 cluster-specific genes were input into Enrichr to determine major cellular subtypes based on gene ontology (enriched biological processes) and cell type (enriched ARCHS4 Tissues) [25, 26].

Cell trajectories were determined using p-Creode (https://github.com/KenLauLab/pCreode) [18]. First, neighborhood variance ratio (NVR) gene selection was used to select genes with local and monotonic variation such that the selected genes possess specific expression patterns over the entire data space amenable to trajectory analysis (https://github.com/KenLauLab/NVR) [27, 28]. Then, cells in two-dimensional expression space were downsampled and a density-based k-nearest neighbor network was constructed. Downsampling was performed with a radius of 30, noise of 8, and target density of 25, which downsampled the original 9,924 input cells to 5,615 (57%). End states were identified by K-means clustering and silhouette scoring of cells with low closeness values (<mean), and the number of end states was doubled to account for rare cell types. A topology was created using a hierarchical placement strategy of cells on path nodes between end states which allowed for the placement of data points along an ancestral continuum. Finally, a representative topology was extracted using p-Creode scoring from an ensemble of n=100 topologies. Graph ID number 3 was chosen based on biological knowledge.

### RNA FISH

Hindlimbs from one pup per litter (time point) were placed in OCT and frozen. Blocks were cryo-sectioned at 10μm thickness and fixed in 4% paraformaldehyde for 15 minutes. Sections were hybridized with probes for mouse collagen, type II, alpha 1 (*Col2a1*), keratin 14 (*Krt14*) and myogenin (*Myog*). A second set was hybridized with probes for collagen, type I, alpha 1 (*Col1a1*), scleraxis (*Scx*), and Runx family transcription factor 2 (*Runx2*) following the manufacturer’s instructions (Advanced Cell Diagnostics, Inc., RNAscope® Fluorescent Multiplex Assay) [29].

### Histology

Hindlimbs from one pup per litter (time point) were fixed in 4% paraformaldehyde for 24 hours and processed for paraffin embedding. Paraffin embedded samples were sectioned at 5μm thickness and stained with Hematoxylin, Safranin-O and Fast Green.

## Acknowledgments

Supported in part by grants from the NIH (AR50245, AR46927, AG15768, AR48182, AR057235, AR073752), Arthritis Foundation, and Nancy Taylor Foundation for Chronic Diseases. We thank the Genome Technology Access Center in the Department of Genetics at Washington University School of Medicine. The authors thank Dr. Samantha Morris for discussions in the early stages of this work.

Authors’ roles: Study design: NHK, NPTH, FG. Study conduct: NHK, NPTH. Data collection: NHK, NPTH. Data analysis: NHK. Data interpretation: NHK, NPTH, FG. Drafting manuscript: NHK. Revising manuscript content: NHK, NPTH, FG. Approving final version of manuscript: all authors. NHK takes responsibility for the integrity of the data analysis.

## SUPPLEMENTARY FIGURES/TABLES

**Supplementary Figure 1:**
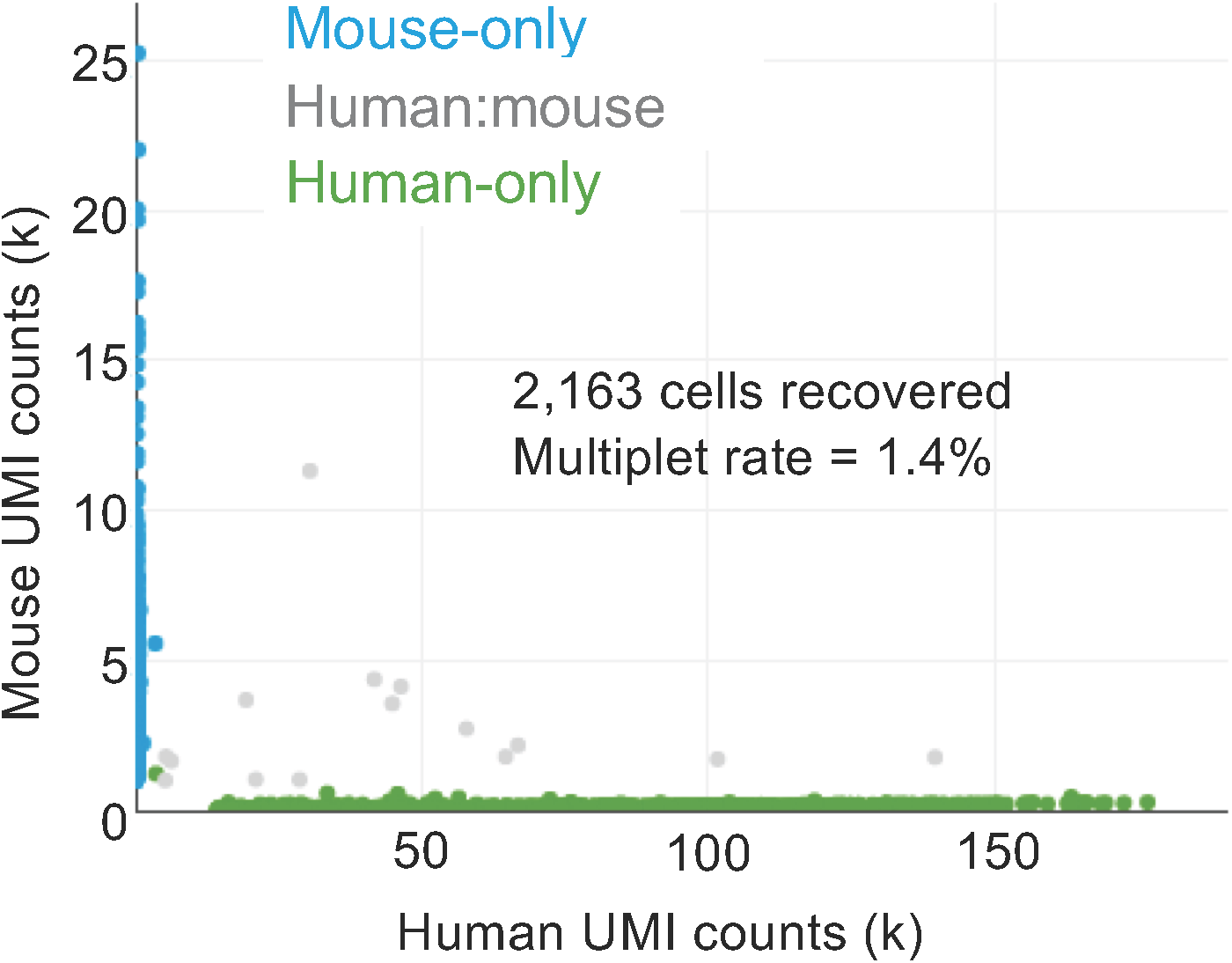
Single cell capture was efficient. Scatter plot of human and mouse UMI counts detected in a mixture of 293T (human) and 3T3 (mouse) cells. “Mouse-only” indicates cell barcodes containing primarily mouse reads (blue); “Human-only” indicates cell barcodes with primarily human reads (green); “Human:mouse” indicates cell barcodes with significant mouse and human reads (grey). A multiplet rate of 1.4% was determined.

**Supplementary Figure 2:**
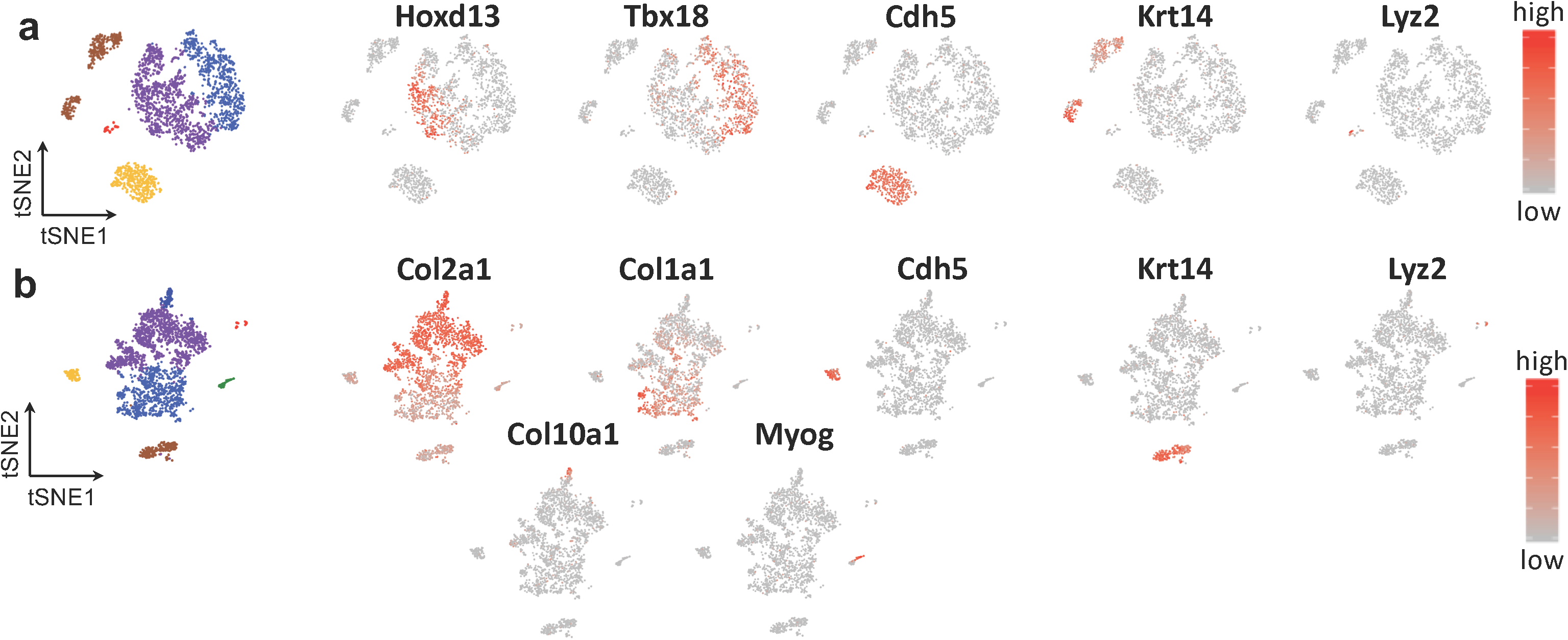
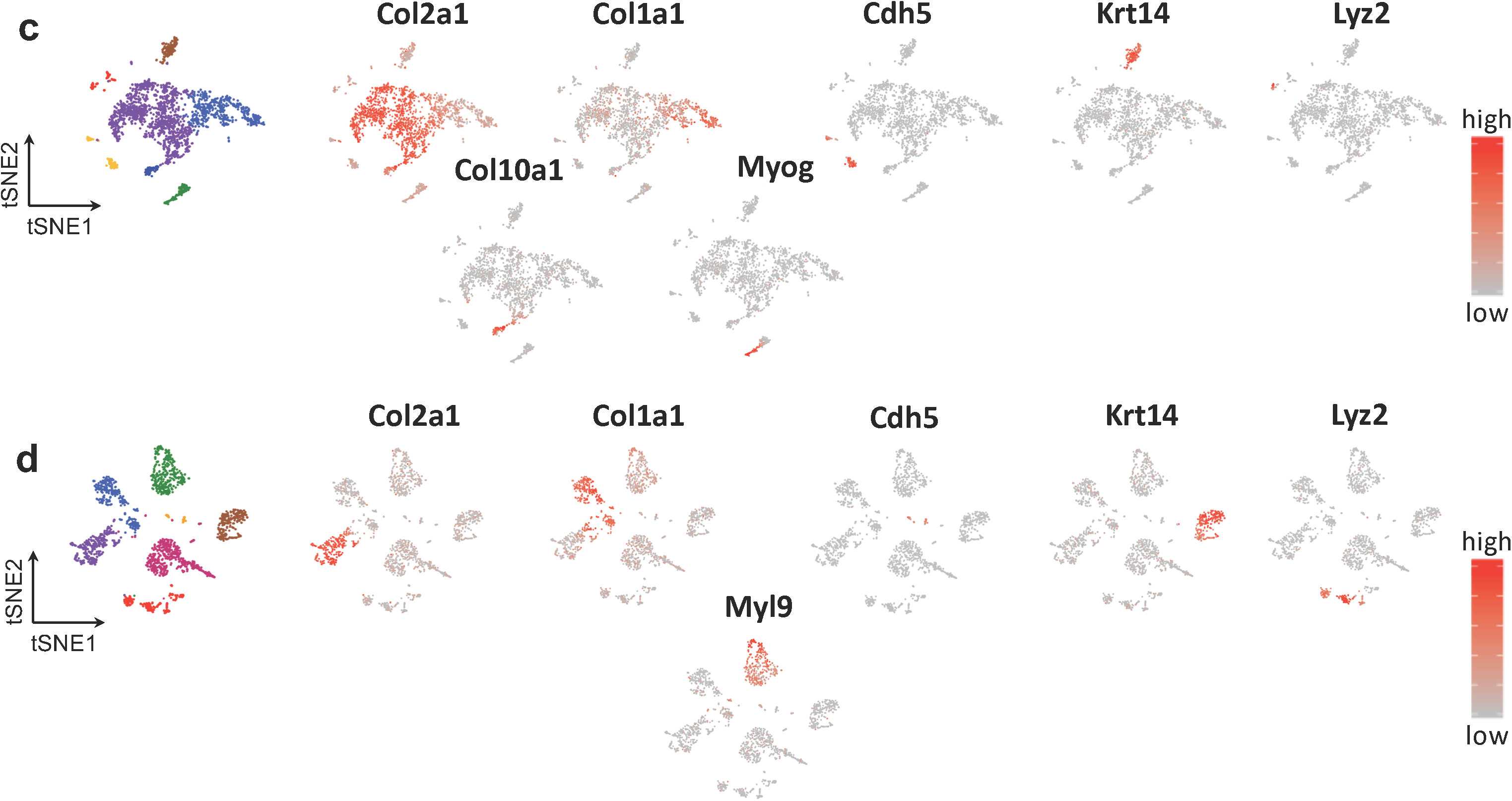
Expression patterns of marker genes at individual time points. Related to Fig. 1. Normalized expression patterns of selected genes to identify the different cell populations in the unsupervised clustering, overlaid on the tSNE projections from samples **(a)** E11.5, **(b)** E13.5, **(c)** E15.5, and **(d)** E18.5.

**Supplementary Figure 3:**
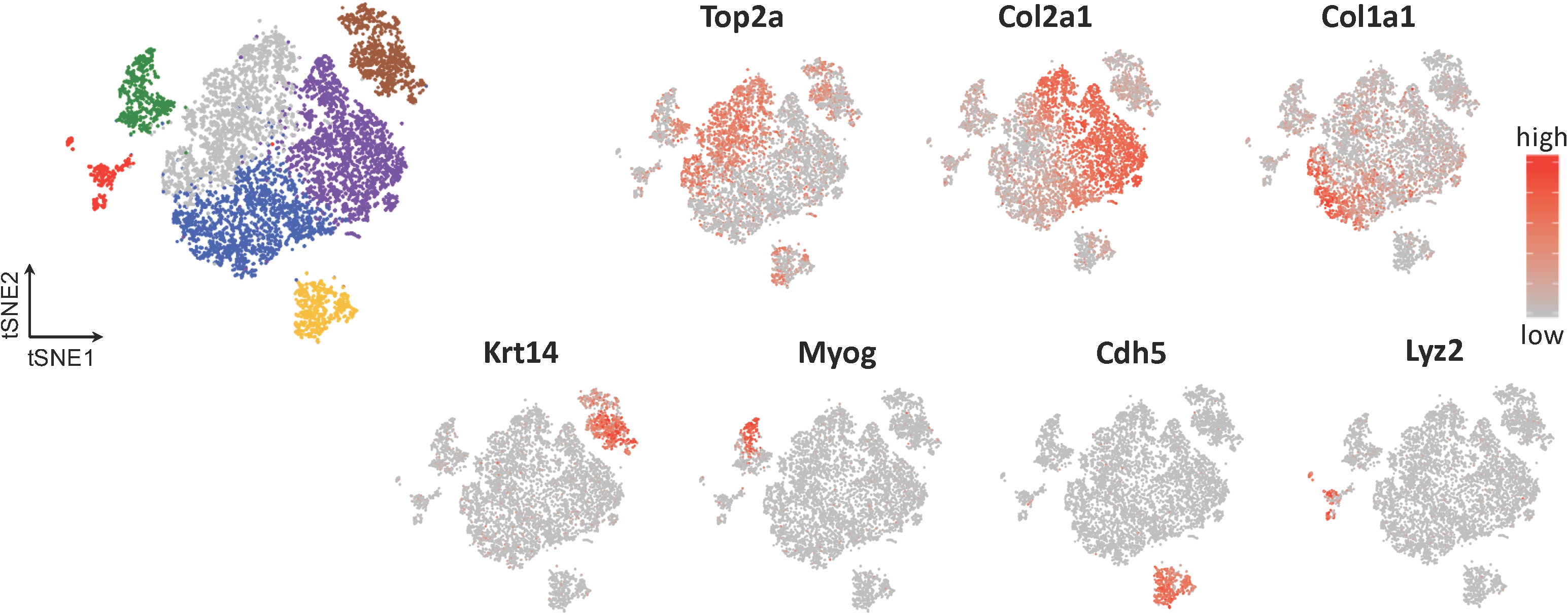
Expression patterns of marker genes across the combined time points. Related to Fig. 2. Normalized expression patterns of selected genes to identify the different cell populations in the unsupervised clustering, overlaid on the tSNE projection from the CCA combined samples.

